# Inflammatory macrophages drive smooth muscle dedifferentiation via YAP signaling in murine deep vein thrombosis

**DOI:** 10.1101/2025.09.24.678378

**Authors:** Huan Yang, Ting Zhou, Jooyong Kim, Bo Liu

**Author notes:** Correspondence to: Bo Liu, PhD. Ting Zhou, PhD. University of Wisconsin-Madison 1111 Highland Avenue, WIMR 4418, Madison, WI 53705.

## Abstract

**Background:** Deep vein thrombosis (DVT) is a common clinical problem characterized by the formation of blood clots in deep veins. The contribution of venous smooth muscle cells (SMCs) to the onset of DVT, particularly the mechanisms driving their phenotypic changes, remains poorly understood.

**Methods:** We employed a murine stasis model of DVT via inferior vena cava (IVC) ligation in both male and female mice. Single-cell RNA sequencing (scRNA-seq) was used to analyze the cellular and transcriptomic landscape of the vein wall 24 hours post-ligation. To investigate cellular crosstalk, we utilized an in vitro system where primary mouse aortic SMCs were treated with conditioned media from inflammatory (M1-like) bone marrow-derived macrophages (BMDMs). The effects on SMC phenotype and signaling were assessed using bulk RNA-seq, Western blotting, qRT-PCR, and functional assays.

**Results:** scRNA-seq analysis revealed that DVT induction promotes a rapid inflammatory response characterized by neutrophil influx and a shift in SMCs from a contractile to a synthetic phenotype in both sexes. Infiltrating myeloid cells were identified as a primary source of signaling to SMCs. In vitro, conditioned media from M1-like macrophages was sufficient to suppress SMC contractile gene expression and function. Mechanistically, this effect was linked to the inhibition of the Hippo pathway effector YAP, evidenced by increased YAP phosphorylation (S127/S397) and a subsequent reduction of total YAP protein in SMCs. We identified macrophage-derived IL-1β as a key ligand that mimics these effects. Importantly, pharmacological activation of YAP rescued contractile gene expression in SMCs exposed to the inflammatory macrophage secretome.

**Conclusions:** Our findings demonstrate that SMC dedifferentiation occurs early in DVT, driven by inflammatory macrophage-mediated suppression of YAP signaling. This study uncovers a critical macrophage-SMC crosstalk mechanism in early DVT pathogenesis and highlights YAP as a potential therapeutic target to preserve vein wall integrity and prevent long-term complications.

## BACKGROUND

Deep vein thrombosis (DVT), the formation of blood clots in deep veins, is a common vascular disease that can lead to severe complications including pulmonary embolism. DVT can also cause post-thrombotic syndrome, a condition characterized by long-term vein wall damage and remodeling [1, 2]. The pathophysiology of DVT involves complex interactions among coagulation factors, blood cells, and the vein wall [3]. While the contribution of the endothelium is well-recognized, the role of smooth muscle cells (SMCs)—another major cellular constituent of the vein wall—remains less understood [3]. Due to limited access to affected vein wall tissue from patients, investigations into venous remodeling rely heavily on animal models, including the full or partial ligation of the inferior vena cava (IVC)—referred to as stasis and stenosis models, respectively— as well as ferric chloride and electrolytic injury models [4, 5]. Using the stasis model of IVC ligation in male mice, our previous single-cell RNA sequencing (scRNA-seq) study characterized the cellular and molecular signatures of the vein wall 24 hours after thrombosis [6]. We observed a dramatic shift in the cellular landscape, dominated by neutrophil infiltration and a reduction in vascular cells. Notably, SMCs displayed clear transcriptomic changes indicative of dedifferentiation, including the downregulation of contractile genes. However, the mechanisms driving this process and whether a similar phenotypic shift occurs in female mice remain unknown.

The contribution of SMC phenotypic change to vascular pathophysiology has been reported in the context of arterial diseases including atherosclerosis, restenosis, and aortic aneurysm [7–9]. Experimental evidence from linage tracing combined with scRNA-seq showed that SMCs can shift from a differentiated contractile phenotype to other functional phenotypes, such as proliferation, migration, inflammation, or fibroblast-like [7, 8]. Multiple transcription factors and co-factors as well as epigenetic mechanisms have been shown to regulate the transition of SMC from a contractile to a de-differentiated state in the context of arterial disease [10, 11]. While the underlying molecular mechanisms may vary depending on disease or the nature of injury, diminished expression of contractile genes including *Acta2*, *Myh11*, and *Cnn1* is a common characteristic of phenotypic shift [7, 8].

In this study, we used scRNA-seq to characterize the SMC phenotypic shift in both male and female mice in the stasis DVT model. In addition, we explored how inflammatory cells influence SMC dedifferentiation using an in vitro cell-cell communication system.

## METHODS

### Animal Studies and DVT Model

All animal procedures were conducted under the approval of the University of Wisconsin—Madison Animal Care and Use Committee (Protocol #M005792). All experiments used 8-to 12-week-old, weight-matched male or female C57BL/6J mice (The Jackson Laboratory, Stock #000664).

The mouse stasis model of DVT was induced through inferior vena cava (IVC) ligation [6]. Mice were anesthetized with 5% isoflurane and maintained on 2.5% isoflurane thereafter. 0.6 mg/kg buprenorphine was administered subcutaneously for pain management before a midline laparotomy was performed. The IVC was exposed, and its back and side branches were respectively cauterized and ligated. The IVC was then fully ligated with 7/0 polypropylene suture just below the left renal vein. For control animals, a sham surgery consisting of vessel dissection without ligation was performed.

### Isolation and Culture of Primary Mouse Aortic Smooth Muscle Cells and Bone Marrow-derived M**φ**s

Primary mouse SMCs were isolated from the aortas of 6-8 weeks old C57BL/6J male mice, as previously described [12]. SMCs were grown to confluence in DMEM (Gibco) supplemented with 10 fetal bovine serum (FBS, Gibco), 100 U/mL penicillin, and 100 µg/mL streptomycin at 37 °C in 5% CO_2_. The experiments were performed with SMCs between passages 2 and 10.

To generate bone marrow-derived macrophages (BMDMs), bone marrow was harvested from the femurs and tibias of mice, as previously described [13]. The cells were cultured in macrophage medium, consisting of DMEM supplemented with 10% FBS, 1% penicillin/streptomycin, and 20 ng/mL of murine M-CSF. Cells were maintained at 37°C and 5% CO_2_, with a complete medium change after three days. Following a total of seven days of differentiation, the adherent cells were ready for subsequent experiments.

For the treatment of SMCs with BMDM conditioned medium, primary SMCs were serum-starved in DMEM containing 0.3% FBS, 100 U/mL penicillin, and 100 µg/mL streptomycin for 48 hours. In the meantime, BMDMs were first primed with 20ng/mL IFNγ overnight, followed by a 4-hour LPS stimulation (100 ng/mL). The IFNγ/LPS treated and control BMDMs were then thoroughly washed with PBS 4 times. Fresh DMEM containing 0.3% FBS, 100 U/mL penicillin, 100 µg/mL streptomycin and 10ng/mL M-CSF were added to the BMDMs. After a 24-hour incubation period, the conditioned media was harvested and centrifuged at 1000 g for 5 minutes to remove any cell debris. Subsequently, the SMCs were exposed to the conditioned media, which was prepared from equal numbers of BMDMs.

### Sample Collection and Preparation

Mice were euthanized 24 hours after surgery. For histological studies, the IVC with the thrombus was collected, embedded, and sectioned for immunofluorescent staining.

For scRNA-seq, IVCs were carefully dissected from adjacent tissues and any intraluminal thrombus, and subjected to a two-step enzymatic digestion to generate a single-cell suspension. Cells from 6 DVT-adjacent IVCs or 6 sham IVCs were pooled to form one sample. 10,000 cells per sample were loaded using the 10x Genomics Chromium platform and the Single Cell 3’ v3.1 Reagent Kit to construct libraries, which were then sequenced on an Illumina NovaSeq X Plus flowcell using a 2×150 bp sequencing reaction targeting >90,000 reads/cell.

For bulk RNA-seq, primary SMCs were treated with conditioned media, harvested, and RNA was extracted. The extracted RNA was then used to synthesize cDNA and prepare a library using an Illumina Stranded mRNA Prep kit, following the manufacturer’s instructions. The cDNA library was then subjected to Illumina NovaSeq X Plus sequencing with the settings of 2×150 bp and 50 million reads per sample.

### Bioinformatic Analysis

For scRNA-seq, raw sequencing reads were aligned to the mouse reference genome (mm10) and quantified using Cell Ranger Count (v6.1.2). The resulting data was analyzed with the OmicVerse package (v1.7.1) [14] on the Python 3.11 platform. After removing low-quality cells, data was normalized and integrated across samples. Cell clusters were identified, and their identities were confirmed by identifying enriched genes and examining the expression of known canonical markers.

Differential expression analysis between the DVT and sham groups was performed using the MAST package, with differentially expressed genes (DEGs) identified based on a log2 fold change > 0.25 and a Bonferroni adjusted p-value < 0.0001. These DEGs were then used for pathway enrichment analysis with clusterProfiler (v4.16). To investigate cellular crosstalk, cell-cell communication networks were inferred using CellChat (v2.1), with a significance threshold of p < 0.05.

For bulk RNA-seq analysis, salmon (version 1.10.1) [15] was employed to quantify transcript levels. DESeq2 was used to normalize the counts and identify differentially expressed genes (DEGs). Subsequently, these DEGs were utilized for pathway enrichment analysis using clusterProfiler. Ligand activities mediating BMDM-SMC intercellular communication were predicted by using the nichenetr package (v2.2.0) [16]. BMDM bulk RNA-seq datasets were downloaded from GEO (GSE103958).

### Immunofluorescent staining

Tissue sections were fixed with 4% PFA, permeabilized with 0.1% Triton X-100, and blocked with 3% donkey serum. Sections were incubated at 4 °C overnight in 0.3% BSA containing primary antibodies: FITC anti-α-smooth muscle actin (ab8211, Abcam; 1:500), anti-Phospho-YAP (Ser127) (13008, Cell Signaling Technology; 1:100), anti-YAP (14074, Cell Signaling Technology, 1:100). After several washes with PBS, sections were incubated with Alexa Fluor 488- or Alexa Fluor 594-conjugated secondary antibodies for 1 h at room temperature. Negative control slides were stained with secondary antibody only. DAPI-containing mounting media (GBI Labs, Catalog #E19-100) was used as a counterstain. Images were acquired with a Nikon A1RS confocal microscope system and analyzed using Fiji software.

### Western Blotting

Cells were lysed in RIPA buffer (R0278, Sigma-Aldrich) containing protease and phosphatase inhibitors (Halt Cocktail, Thermo Scientific). Protein extracts were loaded and separated by SDS-PAGE and then transferred to polyvinylidene fluoride membranes. The membranes were blocked with 5% skim milk, and incubated overnight at 4 °C with the following primary antibodies: anti-Phospho-YAP (Ser127) (13008, Cell Signaling Technology; 1:1000), anti-YAP (14074, Cell Signaling Technology, 1:1000), anti-Phospho-YAP (Ser397) (13619, Cell Signaling Technology; 1:1000), anti-SM22 (36090, Cell Signaling Technology; 1:1000), anti-MYH11 (ab133567, Abcam; 1:1000), anti-Pan-TEAD (13295, Cell Signaling Technology, 1:1000), anti-α-smooth muscle actin (19245, Cell Signaling Technology, 1:1000), anti-β-actin (A5441, Sigma-Aldrich, 1:5000), followed by HRP (horseradish peroxidase)-labeled secondary antibody (Jackson ImmunoResearch, 1:5000).

### RT-qPCR

Total RNA was extracted from cultured cells using Trizol reagent (15596018, ThermoFisher Scientific) according to manufacturer’s protocols. One microgram total RNA was used for the first-strand cDNA synthesis followed by real-time RT-qPCR (polymerase chain reaction). Primer sequences used were: Il6: forward 5′-CCACCAAGAACGATAGTCAATTCC-3′, reverse 5′-GCCATTGCACAACTCTTTTCTCA-3′; Ccl2: forward 5′-ATTCACCAGCAAGATGATCCCAAT-3′, reverse 5′-TGAGCTTGGTGACAAAAACTACAG-3′; Cnn2: forward 5’-ACAAGAGCGGAGATTTGAGCCG-3’, reverse 5’-TCATAGAGGTGACGCCGTGTAC-3’; Myh11: forward 5’-GCAACTACAGGCTGAGAGGAAG-3’, reverse 5’-TCAGCCGTGACCTTCTCTAGCT’-3’; Eln: forward 5’-TCCTGGGATTGGAGGCATTGCA-3’, revers 5’-ACCAGGCACTAAACCTCCAGCA-3’; and Actb: forward 5′-AGCCTTCCTTCTTGGGTATGG-3′, reverse 5′-AAGGGTGTAAAACGCAGCTCA-3′.

### Collagen Gel Contraction Assay

Collagen gel contraction assay was performed as described previously [17]. Briefly, a collagen suspension (1.5 mg/mL) containing trypsin-digested SMCs was cast in one well of a 24-well culture plate. The suspension contained 0.6 ml (1.5×10^5^ cells). The gel was allowed to polymerize for 30 minutes at 37°C. Once polymerized, the gel was carefully detached from the culture well using a pipette tip and immediately exposed to conditioned media at 37°C. The gels were incubated for 24 hours. The collagen gel size change was measured at various times using a ruler and quantified using Fiji image analysis software.

### Scratch Wound Healing Assay

1.5×10^5^ SMCs were seeded into a 24-well plate and allowed to grow until they form a confluent monolayer. The wound was created by using a pipette tip. After 24 hours, the cells were then observed under a microscope and images were captured to measure the extent of wound closure by using Fiji image analysis software.

### Co-immunoprecipitation

Cells were lysed in Pierce IP Lysis Buffer (Thermo Scientific, 87787) and co-immunoprecipitation experiments were performed using SureBeads magnetic beads (Biorad, 1614013) as per the manufacturer’s protocol. In brief, Protein A magnetic beads were washed in TBST, incubated with anti-YAP (14074, Cell Signaling Technology, 1:1000) or rabbit IgG isotype control (Thermo Scientific, 026102) for 30 minutes at room temperature, and then magnetized and washed three times with TBST. The beads were then incubated with cell lysate overnight at 4°C. After incubation, the beads were washed three times with TBST, and the immunoprecipitated proteins were eluted in 1x Laemmli buffer and subjected to pan-TEAD immunoblotting.

### Statistical Analysis

Results are presented as mean ± standard deviation. To assess normality, Shapiro-Wilk normality tests were conducted on the data. If the data did not exhibit a normal distribution, they were log2-transformed and retested for normality. For normally distributed data, a two-tailed Student t test was used for comparison between two conditions. For non-normally distributed data after transformation, a Mann-Whitney U nonparametric test was employed. For comparisons of ≥3 means, a one-way ANOVA with a Tukey post hoc test was used for normally distributed data, while a Kruskal-Wallis nonparametric test was used for non-normally distributed data. For comparisons of how a response is affected by two factors, a two-way ANOVA followed by Sidak multiple comparisons was performed. Statistical analyses were conducted using GraphPad Prism 9.0 (GraphPad Software, Inc). Experiments were repeated as indicated, and differences with p < 0.05 were considered statistically significant.

## RESULTS

### Vein wall cellular landscapes in female and male mouse DVT

We began to investigate SMC phenotypic change during DVT development by performing IVC ligation or sham surgery in age (8 to 12 weeks) matched male and female C57BL/6J mice. 24 hours after the procedure, when blood clots were visibly formed but not yet firmly attached to the vein wall, the IVC tissues were collected, pooled (6 mice per sex) and proceeded to single cell isolation and scRNA-seq. Based on the transcriptomic profiles, cells were separated into different clusters on the Minimum-Distortion Embedding (MDE) plot. At 24-hour post-ligation, the vein wall from both males and females contained 8 distinct cell types: smooth muscle cells (SMCs), fibroblasts (FBs), endothelial cells (ECs), neutrophils (Neus), monocytes/macrophages (Mο/Mφs), T&NK cells, B cells and dendritic cells (DCs) (Fig. 1A). We also detected a very low number of Schwann cells, however, excluded them from further analyses due to their low abundance. Although the relative changes in cell distributions appeared to be slightly different in males and females, the vein wall of both sexes responded to DVT induction with increased percentage of neutrophils but decrease of SMCs, ECs, FBs and Mο/Mφs (Fig. 1B and Supplemental Table S1). Differentially expressed genes (DEGs) and gene ontology (GO) analysis showed that, in both female and male vein walls, DVT induced higher expression of inflammatory response and leukocyte migration associated genes including *Nos2*, *Cd14*, *Tnf*, *Il1*β, *Mmp9*, and *Cxcr2*. In contrast, DVT suppressed the expression of genes associated with extracellular matrix organization and SMC contractility including *Eln*, *Col1a1*, *Col3a1*, *Myh11*, and *Cnn1* (Fig 1C-E).

**Fig. 1.**
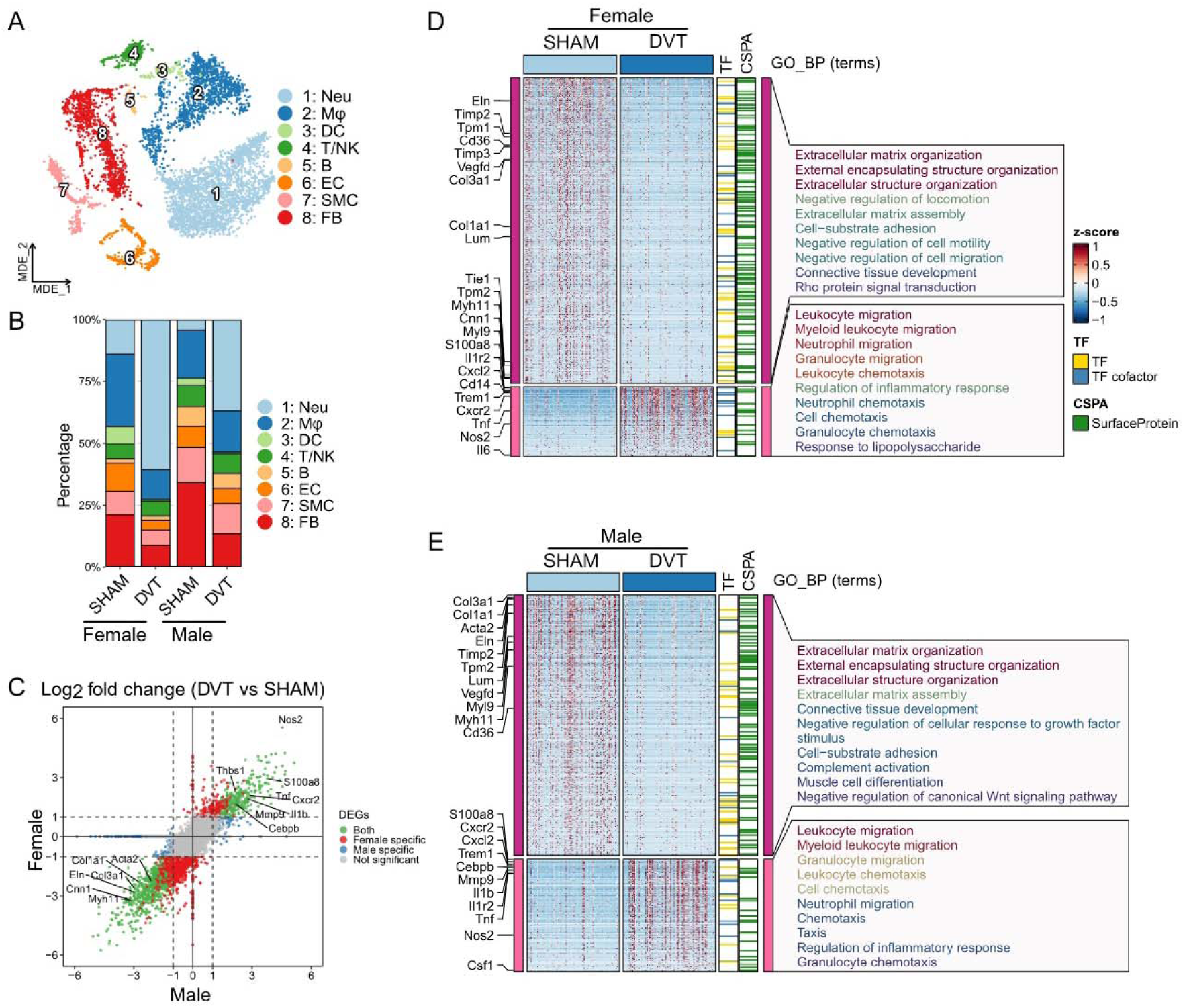
Identification of cell populations in the vein wall of a murine deep vein thrombosis (DVT) model. **(A)** Minimum-Distortion Embedding (MDE) plot of cell clusters identified in the vein wall (male, female, SHAM, and DVT data combined). Neu: neutrophil, DC: dendritic cell, T/NK: T and natural killer cells, B: B cell, EC: endothelial cell, SMC: smooth muscle cell, FB: fibroblast. **(B)** The percentages of cell populations in each group. **(C)** Scatter plot of DVT vs SHAM log2 fold change of gene expression in male and female groups. DEG: differentially expressed gene. **(D&E)** Heatmaps and gene ontology biological process (GO_BP) enrichment analysis of differentially expressed genes in male or female groups (DVT vs SHAM, false discovery rate < 0.05, log2 fold change > 1 or < -1). TF: transcription factor, CSPA: Cell Surface Protein Atlas.

**Supplemental Table S1.**
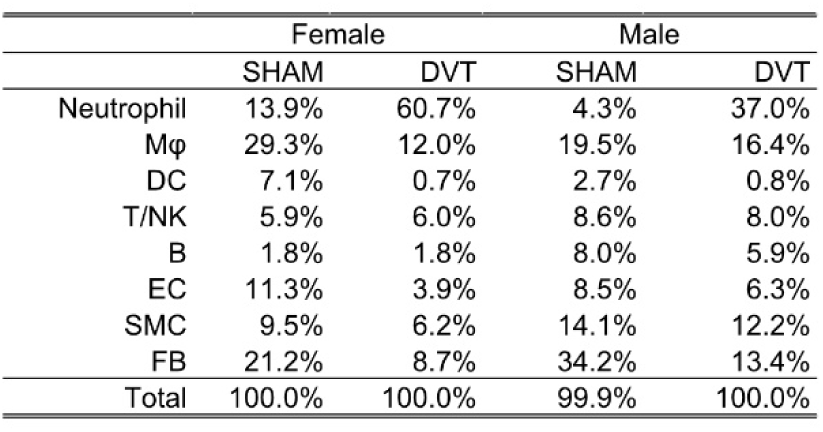
The percentages of cell populations in each group.

|  | Female |  | Male |  |
| --- | --- | --- | --- | --- |
|  | SHAM | DVT | SHAM | DVT |
| Neutrophil | 13.9% | 60.7% | 4.3% | 37.0% |
| Mφ | 29.3% | 12.0% | 19.5% | 16.4% |
| DC | 7.1% | 0.7% | 2.7% | 0.8% |
| T/NK | 5.9% | 6.0% | 8.6% | 8.0% |
| B | 1.8% | 1.8% | 8.0% | 5.9% |
| EC | 11.3% | 3.9% | 8.5% | 6.3% |
| SMC | 9.5% | 6.2% | 14.1% | 12.2% |
| FB | 21.2% | 8.7% | 34.2% | 13.4% |
| Total | 100.0% | 100.0% | 99.9% | 100.0% |

### Cell-cell communications within the vein wall

We next investigated intercellular mechanisms that may govern vein wall remodeling during DVT by analyzing the integrated scRNA-seq dataset using the R package CellChat (Version 2.1). Compared to sham, DVT vein wall showed enhanced intercellular communications from neutrophils and macrophages to vascular cells (ECs, SMCs and FBs) in both sexes, although this effect was more profound in males (Fig. 2A). Similarly, DVT increased EC-to-EC communications in both sexes, with a more significant increase observed in males (Fig. 2A). When examining the signaling pathways from macrophages to SMCs, DVT enhanced signals for SPP1, IL1, GALECTIN, TWEAK, SEMA3, LIGHT, VEGF, and TGFβ in both sexes. In contrast, sham-group veins primarily featured pathways mediated by BMP, IGF, PROS, PTPR, GRN, and GAS (Fig. 2B). A shift in specific signaling pathway was also observed in neutrophil-to-SMC signaling (Fig. 2B). Further examination of the ligand-receptor pairs showed that the communication probabilities of inflammation related ligand-receptor pairs, for example Spp1-Cd44, Il1b-(Il1r1+Il1rap) and Lgals9-P4hb, were enhanced, while anti-inflammation or wound healing associated ligand-receptor interaction such as Grn-Sort1 and Bmp2-(Bmpr1a+Bmpr2) were decreased in DVT vein walls (Fig. 2C).

**Fig. 2.**
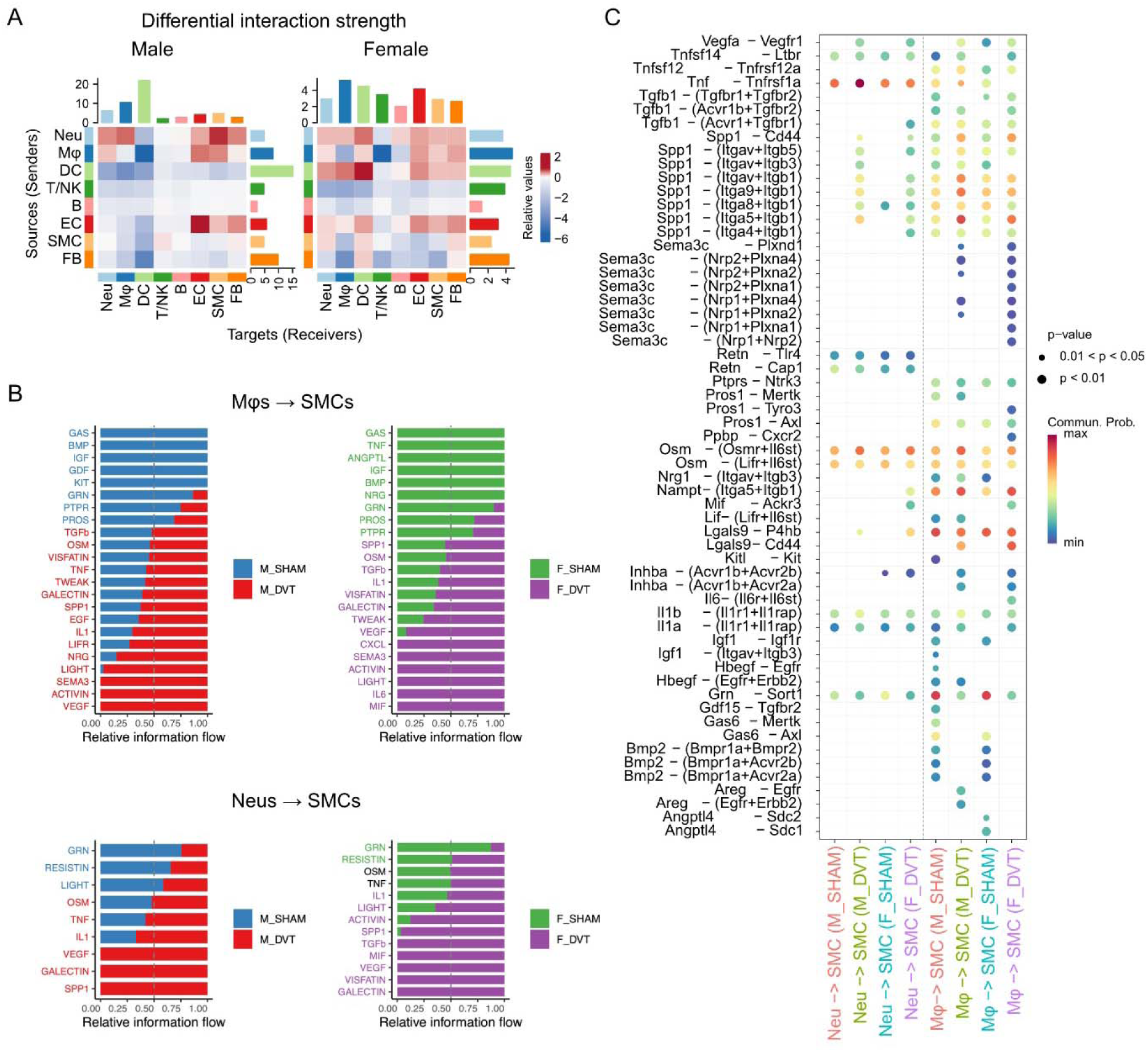
Cell-cell communication in the vein wall. **(A)** Heatmap of differential interaction strength in the DVT group compared to the SHAM group. The top bar plot represents the sum of incoming signals, while the right bar plot represents the sum of outgoing signals. Red (or blue) represents increased (or decreased) signaling in DVT compared to SHAM. Relative value = the interaction strength from source to target in DVT group – the interaction strength from source to target in SHAM group. **(B)** Relative information flow of each signaling pathway from macrophages (Mφs) to smooth muscle cells (SMCs) or from neutrophils (Neus) to SMCs in SHAM and DVT groups. **(C)** Communication probability of ligand-receptor pairs mediating communications between Neus-SMCs or Mφ-SMCs.

### Smooth muscle cell phenotypic switching in DVT

We observed 3 major sub-populations of SMCs in the vein wall of both sexes (Fig. 3A), SMC-1 was enriched in expression of genes related to cell migration, ECM organization and inflammation response like *Il1r1*, *Il1r2*, *Cxcl12* and *Ccl5*, and SMC-3 was likely contractile SMCs which enriched with expressions of muscle structure development associate genes including *Tpm2*, *Acta2* and *Tagln*. SMC-2 was an intermediate state between SMC-1 and SMC-3, with high expression of organelle assembly genes (Fig. 3B). In both sexes, DVT induction altered the distribution of SMC sub-populations by increasing the percentage of SMC-1 while decreasing contractile SMC-3 (Fig. 3C). Trajectory (Fig. 3A), RNA velocity (Fig. 3D) and pseudotime (Fig. 3E) analyses showed that DVT caused a transition from SMC-3 toward SMC-1 in the vein wall. Predicted dynamic gene expression changes during the transition revealed a reduction in smooth muscle contractile genes (*Acta2*, *Cnn1*, *Tagln* and *Tpm2* etc.) but an increase in stimulus response genes (*S100a10*, *Egfr* and *Il1r1*). Gene ontology analysis of these altered genes showed that pathways such as “Muscle system process” and “Muscle contraction” were repressed, however, “Response to growth factor”, “Response to endogenous stimulus” and “Cell migration” pathways were induced along the evolution trajectory (Fig. 3F).

**Fig. 3.**
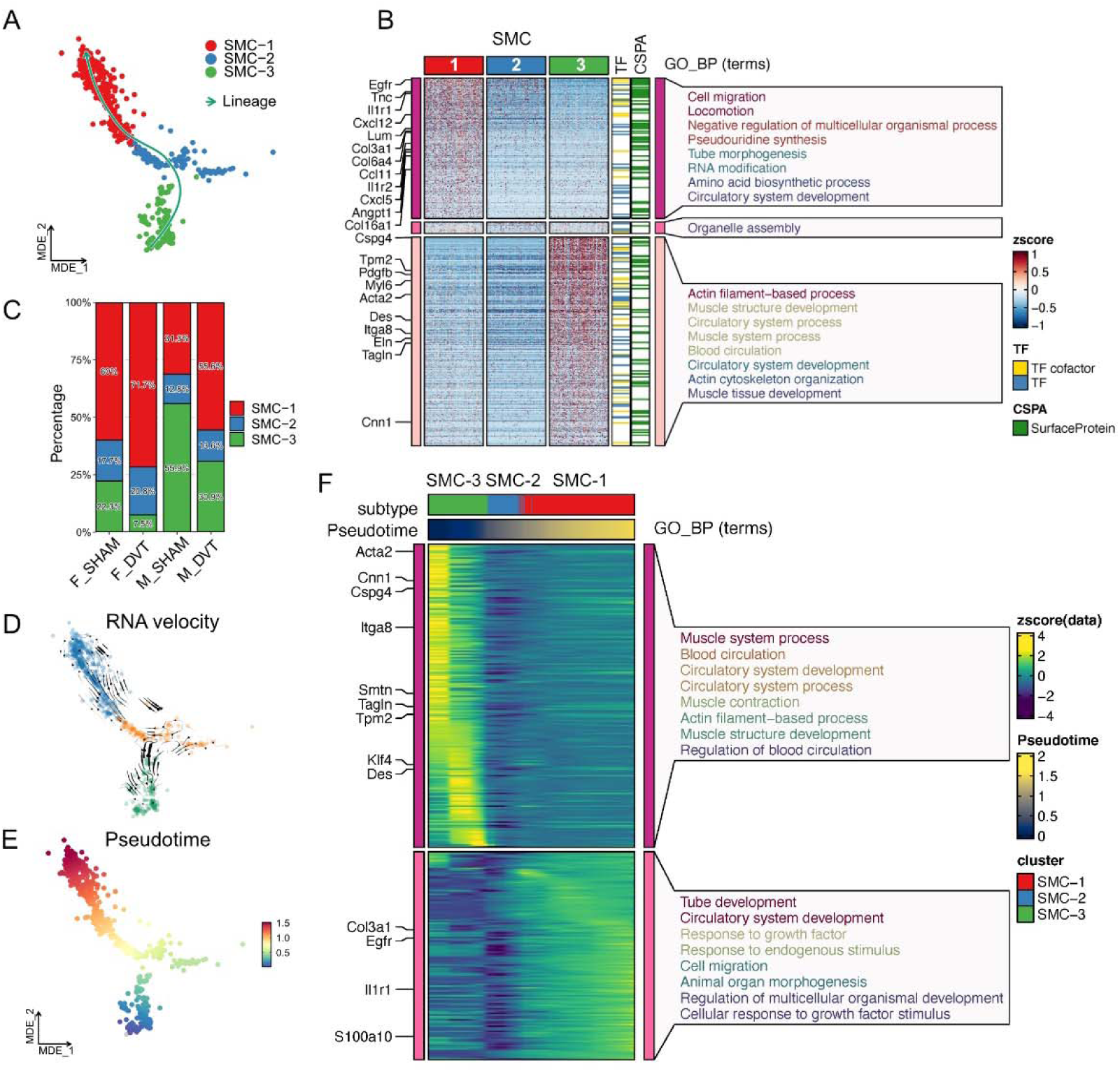
Phenotypic switching of SMCs during DVT. **(A)** Subtypes of SMCs in the vein wall. The green line shows the inferred directed trajectory of cell transition. **(B)** Heatmap of molecular and GO_BP signatures for each SMC sub-cluster. **(C)** The percentages of SMC sub-clusters in SHAM or DVT groups. **(D)** RNA velocity analysis of SMC sub-clusters. The arrow’s direction indicates the predicted future state of a cell, while its length reflects the speed of transcriptional change. **(E)** Pseudotime predicted based on RNA velocity analysis. **(F)** Gene expression changes along the pseudotime axis.

### Population of M1-like pro-inflammatory macrophage expanded in DVT

Having observed enhanced Mφ-to-SMC communication in DVT, we further analyzed macrophage populations in the scRNA-seq data. As shown in Supplemental Figure S1, three Mφ sub-populations were noted. Mφ-1 was characterized by high expression of gene related to regulation of immune response, likely pro-inflammatory or M1-like Mφs; Mφ-3 enriched with genes related to endocytosis and locomotion, likely anti-inflammatory or M2-like Mφs; Mφ-2 was an intermediate stage between Mφ-1 and Mφ-3. In both males and females, DVT increased the expression of pro-inflammatory genes and the percentage of Mφ-1 compared to the sham group, although this pro-inflammatory shift was more pronounced in the vein wall of male mice.

**Supplemental Figure S1.**
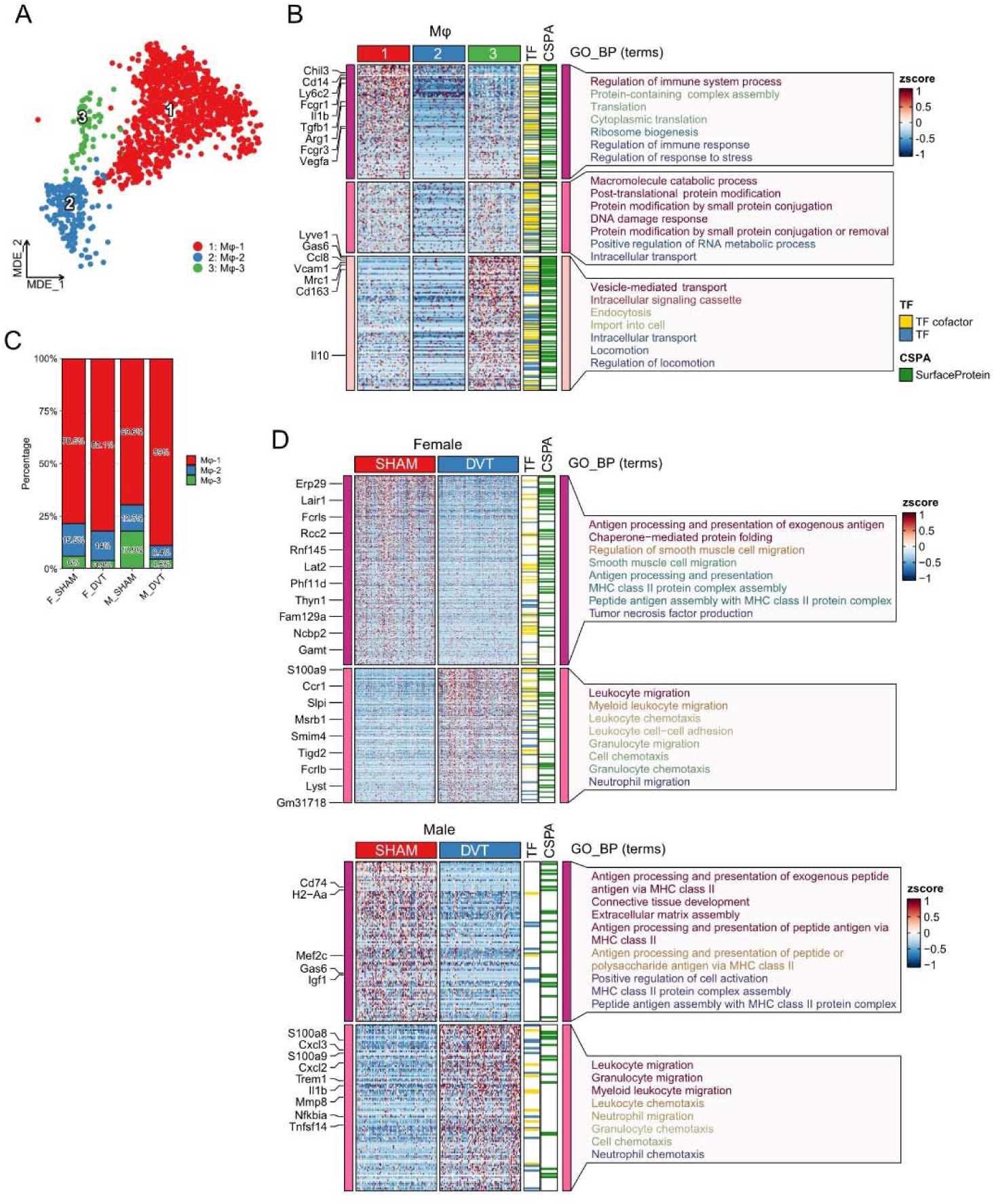
Macrophage populations in DVT. **(A)** Minimum-Distortion Embedding (MDE) plot of macrophage populations in scRNA-seq data. **(B)** The percentages of macrophage populations in SHAM or DVT groups. **(C)** Heatmap of molecular and gene ontology biological process (GO_BP) signatures for each macrophage population. **(D)** Heatmaps and GO_BP enrichment analysis of differentially expressed genes in male or female groups (DVT vs SHAM, false discovery rate < 0.05, log2 fold change > 1 or < -1). TF: transcription factor, CSPA: Cell Surface Protein Atlas.

### Macrophage-SMC communications in vitro

To investigate the signaling mechanism underlying Mφ-to-SMC communication, we conducted bulk RNA sequencing of SMCs after treating them with conditioned media from naïve Mφs (M0-like) or inflammatory M1-like Mφs (polarized with LPS+IFNγ). Principal component analysis (PCA) showed that the transcriptomic profiles of SMCs treated with M1-like conditioned media (referred to as M1-CM) were drastically different from those of SMCs treated with M0-CM or SMCs cultured in regular growth medium (referred to as Reg) (Fig. 4A). DGE analysis showed 1423 up-regulated and 1427 down-regulated genes in M1-CM group (threshold: p < 0.01, log2 fold change > 1 or < -1) compared to M0 group (Fig. 4B). KEGG gene set enrichment analysis (GSEA) suggested that the up-regulated genes were mostly enriched in inflammatory response pathways (representative genes: *Ccl2*, *Il6*, *Nos2* and *Lgals3*) such as TNF signaling pathway, Toll-like receptor signaling pathway, NF-κB signaling pathway, chemokine signaling pathway, as well as apoptosis pathway (*Fas*, *Tnfsf10*, *Casp7*, *Casp10* and *Ripk1*) while down-regulated genes were related to Cardiac muscle contraction (*Myh11*, *Smtn*, *Acta2*, *Tpm1* and *Tpm2*), Cell cycle (*Cdk1*, *Cdc7*, *Mki67* and *Pcna*), Wnt, and Hippo signaling pathway (Fig. 4B&C). Moreover, significant reductions of protein level of MYH11 and mRNA levels of *Cnn1*, *Eln* and *Myh11* in M1-CM group were validated by western blotting and qRT-PCR, respectively (Fig. 4D-G&J), which was consistent with our scRNA-seq analysis showing decreased contractile SMC population (SMC-3) in the DVT vein wall (Fig. 3C). qRT-PCR of *Ccl2* and *Il6* also confirmed the stimulation of inflammation in SMCs treated with M1-CM (Fig. 4H&I).

**Fig. 4.**
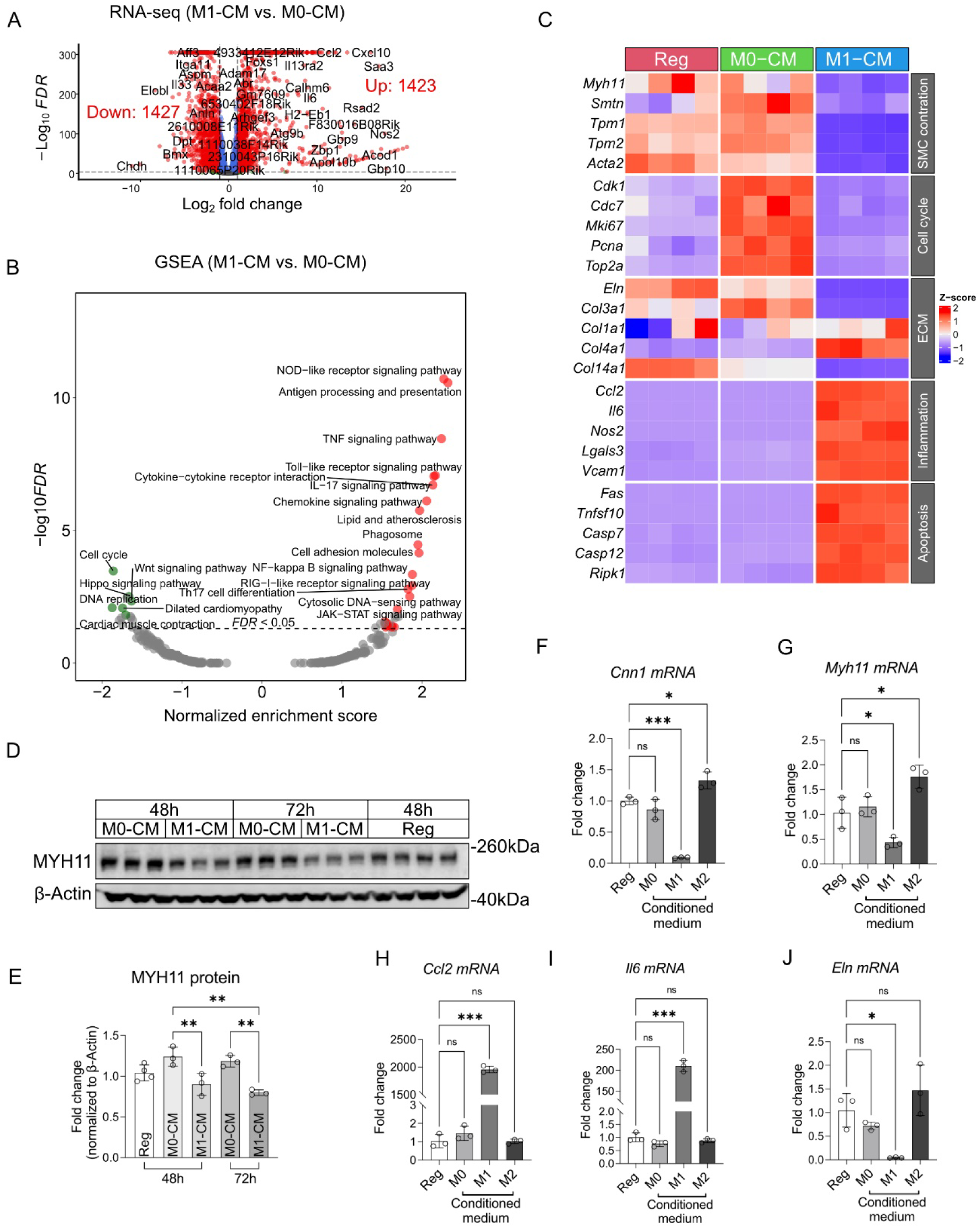
Macrophage-SMC communications in vitro. **(A)** Volcano plot of differentially expressed genes (DEGs) of bulk RNA-seq of mouse primary SMCs treated with indicated conditioned media from bone marrow-derived Mφs (BMDMs) for 24h. **(B)** Gene set enrichment analysis (GSEA) of DEGs using KEGG pathway terms. **(C)** Heatmap showing the expression of representative genes from dysregulated pathways. **(D)** Western blot of MYH11 in mouse primary SMCs treated with indicated conditioned media. **(E)** Quantification of the western blot in (D). **(F-J)** RT-qPCR validation of differentially expressed genes identified by bulk RNA-seq. Data in (E-J) are presented as mean ± SD. One-way ANOVA is performed. \**p* < 0.05, \*\**p*<0.01, \*\*\**p*<0.001.

### Functional analyses of CM-treated SMCs

To determine whether Mφs alter the functions of SMCs, we first measured contractibility using collagen gel contraction assay. M1-CM treatment significantly reduced SMC’s ability to contract, reflected by their diminished ability to shrink the collagen gel (Fig. 5A). We next assessed SMC migration using the scratch assay. As shown Fig. 5B, M0-CM significantly promoted scratch wound closure. M1-CM slightly hindered SMC migration in the scratch assay, albeit insignificantly.

**Fig. 5.**
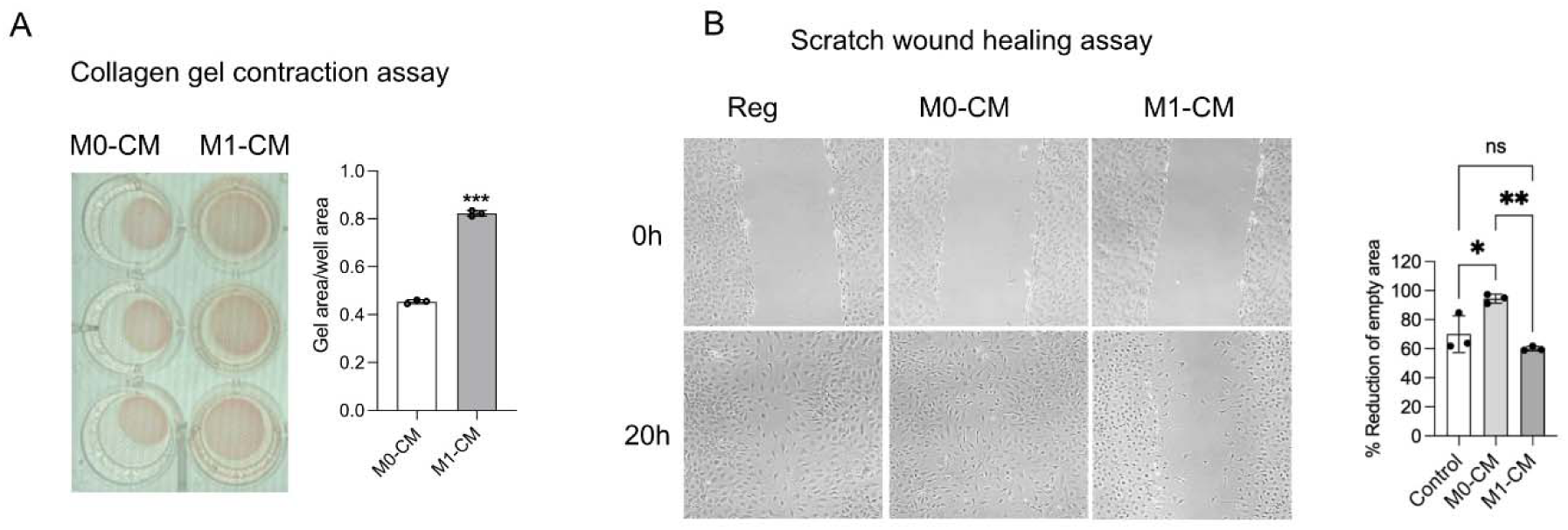
Functional analyses of SMCs treated with BMDM-conditioned media. **(A)** Representative images and quantification of collagen contraction assay of primary mouse SMCs treated with BMDM conditioned media for 24h. **(B)** Representative images and quantification of scratch wound healing assay of primary SMCs treated with BMDM conditioned media. Data in are presented as mean ± SD. Student t test is performed in (A). One-way ANOVA is performed in (B). \**p* < 0.05, \*\**p*<0.01, \*\*\**p*<0.001.

### M1-CM decreased YAP signaling in SMCs

As bulk RNA-seq revealed M1-CM reduced the Hippo signaling pathway in SMCs (Fig. 4B), we examined Yes-associated protein (YAP), the central effector of the Hippo signaling pathway, focusing on YAP phosphorylation (S127 and S397) in CM-treated SMCs. M1-CM increased phosphorylation of YAP S127 and S397, peaked in 3-12 hours and 1-6 hours, respectively (Fig. 6A-C). After 24 hours of CM treatment, total YAP level was significantly lower in the M1-CM group compared to the M0-CM group (Fig. 6D). Since M1-CM did not change the mRNA level of *Yap1* (Fig. 6E), we hypothesized that the reduced abundance of total YAP was due to hyper-phosphorylation and subsequent protein degradation. Consistent with the decreased total YAP level, YAP-TEAD target genes (*Amot*, *Tgfb2*, *Tgfb3*, *Ccn1*, *Ccn4* and *Bmp4*) were also reduced in M1-CM group (Fig. 6E). Notably, M1-CM altered the mRNA expression of YAP binding partners, specifically reducing *Tead2-4* expression but increasing *Tead1* (Fig. 6E). Co-IP assay also showed reduced YAP-TEAD interaction (Fig. 6F), suggesting YAP-TEAD transcription activity was inhibited in M1-CM treated SMCs. To further examined whether YAP mediates the downregulation of contractile genes by M1-CM, we applied PY-60, a small molecule that activates YAP transcriptional activity [18], to SMCs treated with M0-or M1-CM. qRT-PCR result demonstrated that PY-60 robustly promoted *Cnn1* gene expression in both M0-and M1-CM groups (Fig. 6G).

**Fig. 6.**
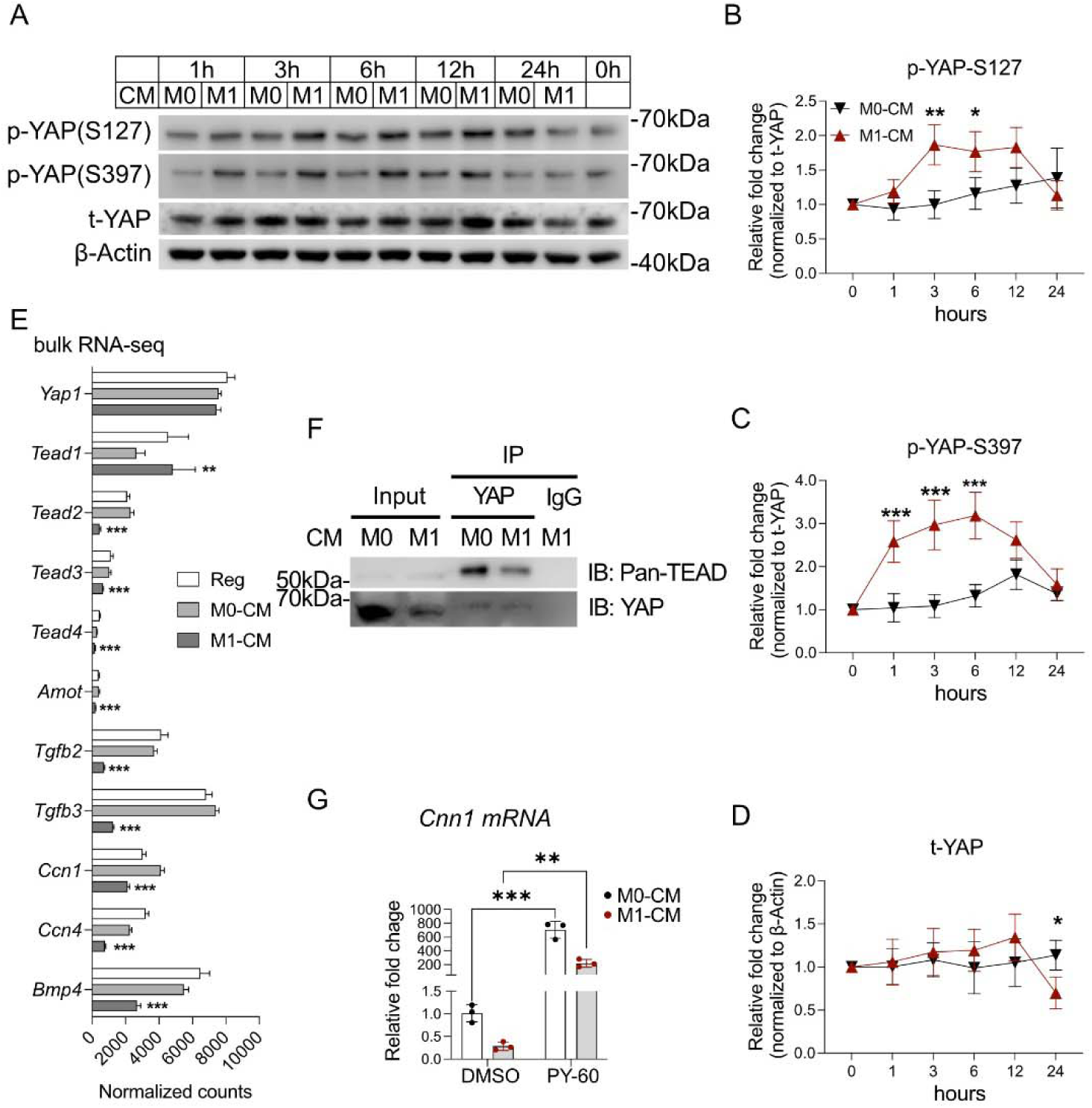
YAP signaling in macrophage conditioned media-treated SMCs. **(A-D)** Time course of phospho-YAP (S127 or S397) and total YAP in mouse primary SMCs treated with M0 or M1-like BMDM conditioned medium (CM). **(E)** Normalized counts of genes associated with YAP-TEAD pathway in SMCs from bulk RNA-seq data. **(F)** Co-immunoprecipitation (Co-IP) of YAP and Pan-TEAD in SMCs treated with M0 or M1-like BMDM conditioned medium. **(G)** RT-qPCR of *Cnn1* in SMCs treated with BMDM-CM or YAP-TEAD pathway activator PY-60 for 24 hours. Data in are presented as mean ± SD. Two-way ANOVA is performed in (B-D) & (G). \**p* < 0.05, \*\**p*<0.01, \*\*\**p*<0.001.

### DVT decreased YAP signaling in the vein wall

To evaluate whether YAP signaling was altered by DVT, we analyzed the vein walls 24 hours after stasis DVT induction or sham surgery by Western blotting. We found significant reduction in phosphor-YAP (S127), total YAP, and pan-TEAD in the DVT group, along with markedly diminished levels of contractile protein MYH11 and α-SMA (Fig. 7A&B). Immunostaining of vein wall cross-sections confirmed the reduction of α-SMA, phospho-and total YAP in the DVT group (Fig. 7C&D). Moreover, YAP was predominantly decreased in venous SMCs (Fig. 7C). Consistently, scRNA-seq analysis showed that SMCs expressed highest level of *Yap1*— along with *Tead1* and *Tead3* — among the major cell types in the vein wall. DVT induction did not change *Yap1* gene expression in SMCs (Fig. 7E), suggesting that transcriptional regulation is unlikely to be the mechanism responsible for the decreased YAP abundance in DVT-affected SMCs.

**Fig. 7.**
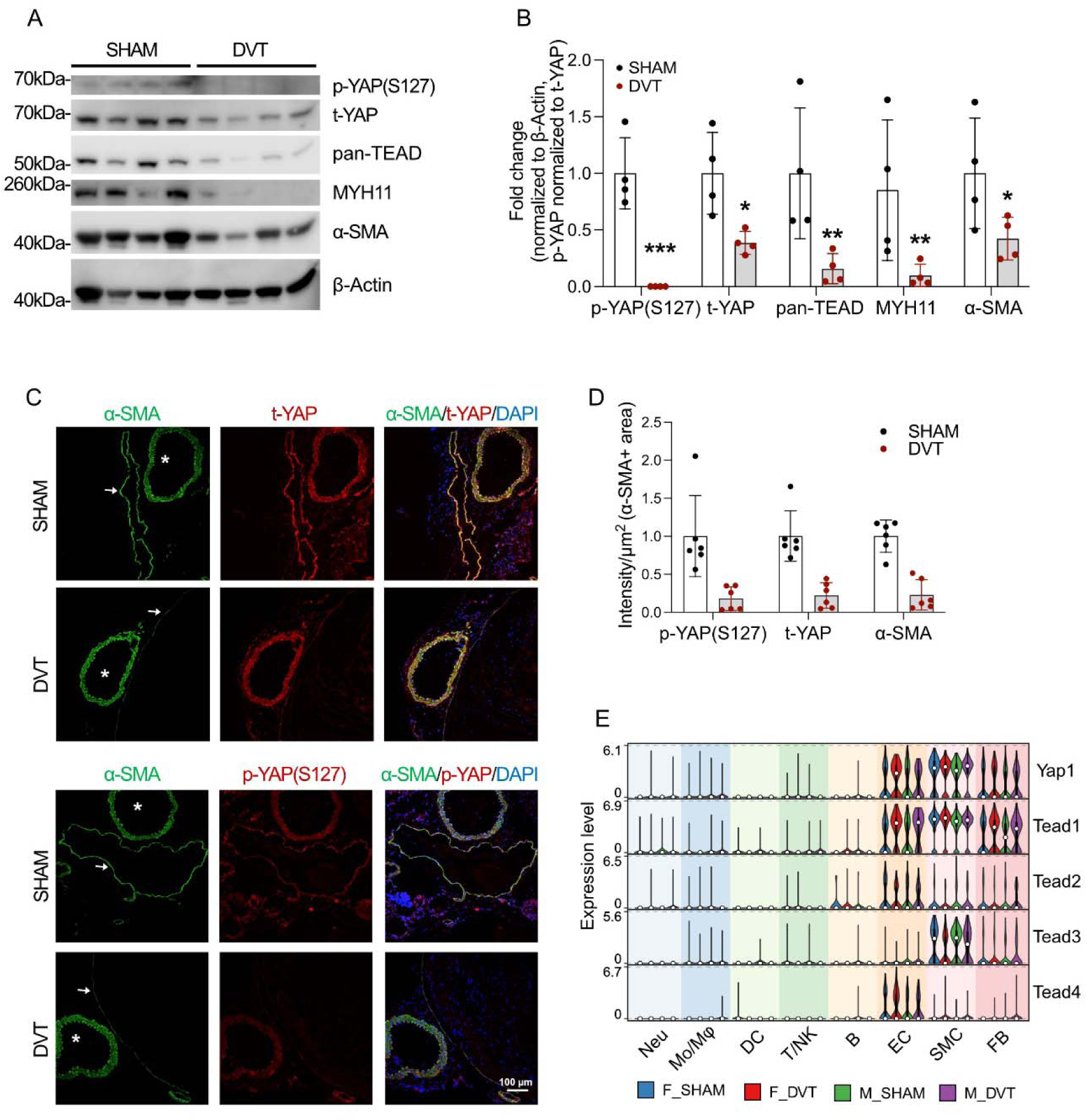
YAP signaling in the vein wall during DVT. **(A-B)** Western blot and quantification of phospho-YAP(S127), total-YAP, pan-TEAD, MYH11 and α-SMA in vein walls 24 hours after SHAM or DVT surgery. **(C-D)** Representative images and quantification of phospho-YAP(S127), total-YAP, and α-SMA in vein walls 24 hours after SHAM or DVT surgery. The white arrow indicates the vein wall; the asterisk indicates the nearby aorta. Scale bar=100 µm. **(E)** *Yap1* and *Tead1-4* expression and distribution in scRNA-seq data. Data in are presented as mean ± SD. Two-way ANOVA is performed in (B&D). \**p* < 0.05, \*\**p*<0.01, \*\*\**p*<0.001.

### M**φ**s communicate with SMCs via IL-1**β**

To identify the ligand-receptor pair(s) that mediate the communication between Mφs and SMCs, we conducted a ligand activity analysis using the R package NicheNet, applying it to both the RNAseq data generated in Fig. 4, and publicly available RNAseq data of BMDM treated IFN-γ and LPS (Das A. et al., GSE103958 [19]). Our analysis predicted that BMDM-expressed IL-1β and TNF had the highest potential to regulate the Hippo signaling pathway in SMCs (Fig. 8A). Interestingly, in the vein wall, *Il1b* was highly expressed by Mφs, while *Il1r1* (the major receptor for IL1β) was enriched in SMCs. The expression levels of *Il1b* and *Il1r1* were elevated by DVT in both sexes (Fig. 8B). To test the role of IL-1β in SMC phenotypic switching, we treated mouse primary SMCs with recombinant mouse IL-1β for various time points. We found that IL-1β significantly increased YAP phosphorylation (S127 and S397) as early as 30 minutes after treatment (Fig. 8C). Prolonged treatment (48h) reduced total YAP, as well as the contractile proteins SM22 and MYH11 (Fig. 8C). The YAP activator PY-60 significantly increased the expression of *Cnn1* and *Myh11* with and without the presence of IL-1β (Fig. 8D).

**Fig. 8.**
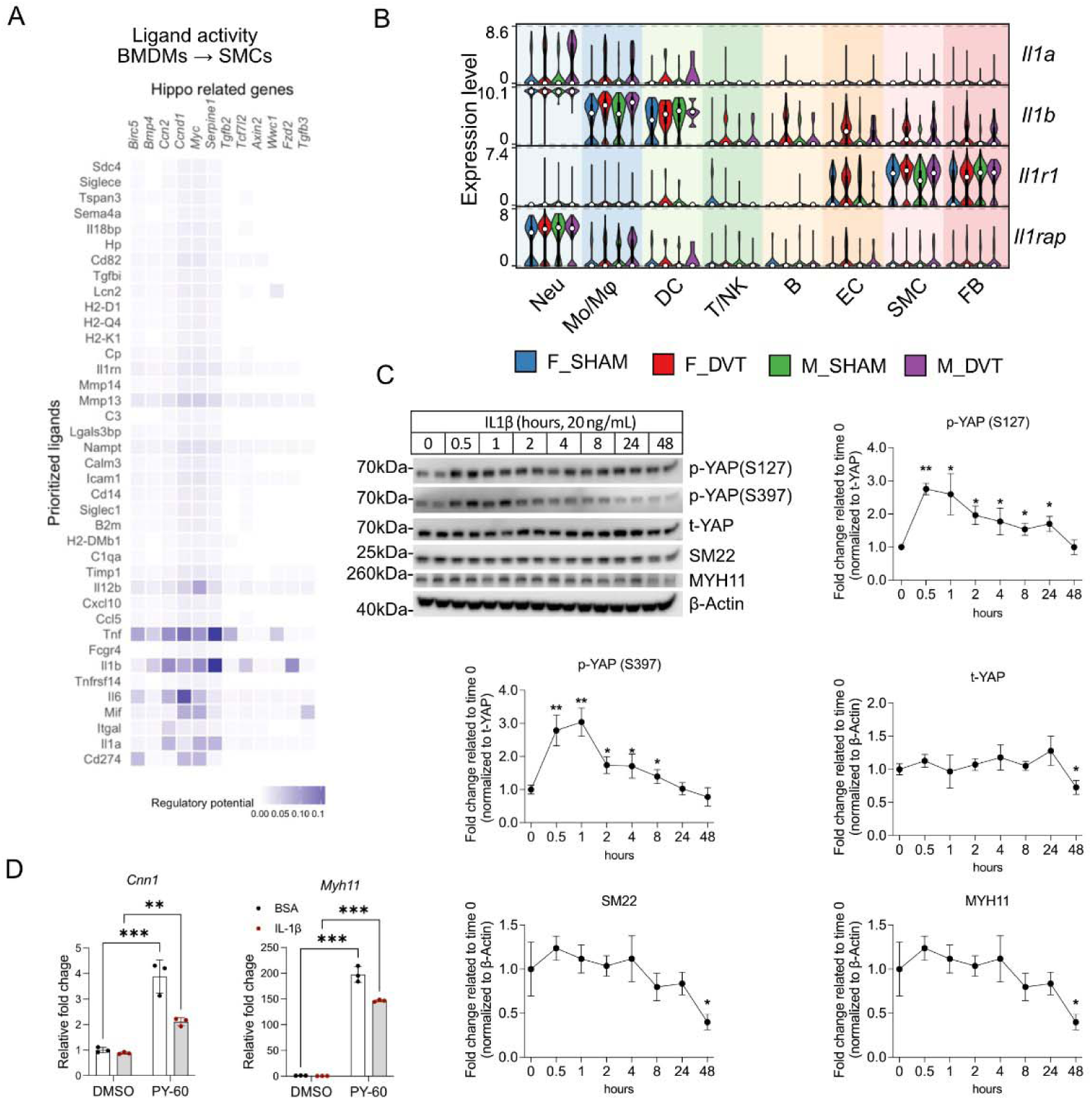
IL-1β mediates macrophage-SMC communication. **(A)** Predicted ligand activities mediating BMDM-SMC communication using the NicheNet R package. IFNγ/LPS stimulated BMDM bulk RNA-seq datasets were downloaded from GEO (GSE103958). **(B)** Expression and cellular distribution of *Il1a*, *Il1b*, *Il1r1* and *Il1rap* in scRNA-seq data. **(C)** Representative Western blot and quantification of the time course of p-YAP(S127), p-YAP(S397), total-YAP, SM22, and MYH11 protein levels in SMCs challenged with 20 ng/mL recombinant mouse IL-1β. **(D)** RT-qPCR of *Ccl2* and *Myh11* in SMCs treated with recombinant mouse IL-1β and/or PY-60. Data in are presented as mean ± SD. Two-way ANOVA is performed in (C&D). \**p* < 0.05, \*\**p*<0.01, \*\*\**p*<0.001.

## DISCUSSION

Our unbiased scRNA-seq analysis revealed profound transcriptomic changes in the mouse vein wall 24 hours after DVT induction. In addition to the expected neutrophil influx, the thrombus-bearing vein wall exhibited phenotypic switching of SMCs, characterized by diminished expression of contractile genes. Another major early DVT response is the shifting of macrophages toward an inflammatory phenotype. SMC fibrosis is believed to contribute to the thickening and stiffening of the vein wall associated with post-thrombotic syndrome [20, 21], which typically occurs several weeks after DVT induction. Our observation of reduced contractile SMCs in both sexes just 24 hours into DVT suggests that SMC phenotypic change begins in the acute phase. This finding points to the potential for early medical interventions to prevent post-thrombotic syndrome.

Several factors released by the luminal thrombi, including PDGF are capable of promoting SMCs switch from the contractile state to synthetic/mesenchymal states. Our scRNA-seq and CellChat analysis identified neutrophils and macrophages that infiltrated the vein wall as novel drivers of SMC phenotypic switching. In full support of this notion, the secretome of inflammatory macrophages drastically altered the gene expression as well as the functions of SMCs in vitro. The bioinformatic analysis of gene expression patterns of SMCs led us to the Hippo signaling pathway. The role of YAP, the central effector of the Hippo signaling, in SMCs has been reported in several arterial diseases. YAP expression was reduced in human aortic aneurysms [22] but increased in vein graft disease [23]. In mouse carotid artery ligation and rat carotid artery balloon injury models, YAP expression was elevated in SMCs, whereas phosphorylated YAP (S127)/total YAP ratio was decreased, in parallel with reduced expression of contractile SMC markers [24]. However, to the best of our knowledge, this study is the first to report that DVT induction in mice increased YAP phosphorylation in the vein wall. Our in vitro data showed that inflammatory macrophages increased YAP phosphorylation and decreased total YAP protein levels in SMCs. Activation of YAP in SMCs preserved contractile gene expression in macrophage-conditioned SMCs, supporting the pro-differentiation role of YAP in SMCs in the context of inflammation. However, whether manipulation of YAP can prevent SMC phenotypic change in DVT and subsequently prevent the fibrotic remodeling of the vein wall warrants further investigation.

Clinically, DVT affects both men and women [25]. In mice, IVC ligation causes intraluminal clotting at a similar rate [5]. Consistently, we found in the mouse model, DVT induction triggered generally similar cellular changes in both sexes such as influx of neutrophils and phenotypic switching of SMCs (toward dedifferentiation) and macrophages (toward inflammation). However, subtle sex-dependent responses were noted. For example, the DVT-triggered increase in the expression of pro-inflammatory genes as well as the presence of inflammatory Mφ macrophage subtype is more pronounced in the vein wall of male mice. Future studies are necessary to address this potential sex dimorphism in DVT.

There are several limitations associated with the current study. First, our study relies on animal and cell modeling due to the lack of access to tissue from human DVT-affected veins, as DVT is rarely treated with open surgery. Second, the transcriptomic focused scRNA-seq and bulk RNAseq analyses do not provide direct evidence of proteomic changes particularly at the level of post-translational modifications. Finally, the specific components of the macrophage secretome responsible for altering SMC phenotype remain to be fully identified. Although IL1β is capable of influencing SMCs, multiple factors from macrophages, neutrophils and platelets are likely to be the collective drivers of SMC phenotypic changes.

## CONCLUSIONS

In this study, we uncovered that SMC phenotypic changes begin during the early phase of murine DVT. The diminished YAP signaling in the vein wall is likely responsible for the reduced contractile gene expression by SMCs within the clot-bearing vein wall. Our work highlights a new role of inflammation in post-thrombotic vein remodeling, which could offer a new treatment direction for DVT patients.

## DECLARATIONS

### Ethics approval and consent to participate

Not applicable

### Consent for publication

Not applicable

### Availability of data and materials

All data and materials are available from the corresponding author on reasonable request. Sequencing data will be made publicly available at the Gene Expression Omnibus upon publication.

### Authors’ contributions

HY conducted the bioinformatic analyses and performed all in vitro experiments. TZ performed animal surgeries and prepared the libraries for single-cell and bulk RNA sequencing. JK assisted with the statistical analyses. HY, TZ, and BL drafted the manuscript. All authors read and approved the final version.

## Acknowledgements

Single-cell RNA sequencing and bulk RNA sequencing was performed at UW Biotechnology Center Gene Expression Center and DNA Sequencing Facility (Research Resource Identifier – RRID:SCR_017757 and SCR_017759).

## List of abbreviations

BMDMs: Bone marrow-derived macrophages
DCs: Dendritic cells
DEGs: Differentially expressed genes
DVT: Deep vein thrombosis
ECs: Endothelial cells
FBs: Fibroblasts
GO: Gene ontology
GSEA: Gene set enrichment analysis
IVC: Inferior vena cava
M1-CM: M1-like conditioned media
Mο/Mφs: Monocytes/macrophages
Neus: Neutrophils
PCA: Principal component analysis
Reg: Regular medium
qRT-PCR: Real-time quantitative polymerase chain reaction
scRNA-seq: Single-cell RNA sequencing
SMCs: Smooth muscle cells
YAP: Yes-associated protein

## REFERENCES

1. Heit JA, Silverstein MD, Mohr DN, Petterson TM, Lohse CM, O’Fallon WM, Melton LJ, 3rd: The epidemiology of venous thromboembolism in the community. Thromb Haemost 2001, 86:452–463.

2. Henke PK, Comerota AJ: An update on etiology, prevention, and therapy of postthrombotic syndrome. J Vasc Surg 2011, 53:500–509.

3. Navarrete S, Solar C, Tapia R, Pereira J, Fuentes E, Palomo I: Pathophysiology of deep vein thrombosis. Clin Exp Med 2023, 23:645–654.

4. Diaz JA, Obi AT, Myers DD, Jr., Wrobleski SK, Henke PK, Mackman N, Wakefield TW: Critical review of mouse models of venous thrombosis. Arterioscler Thromb Vasc Biol 2012, 32:556–562.

5. Diaz JA, Saha P, Cooley B, Palmer OR, Grover SP, Mackman N, Wakefield TW, Henke PK, Smith A, Lal BK: Choosing a Mouse Model of Venous Thrombosis. Arterioscler Thromb Vasc Biol 2019, 39:311–318.

6. DeRoo E, Zhou T, Yang H, Stranz A, Henke P, Liu B: A vein wall cell atlas of murine venous thrombosis determined by single-cell RNA sequencing. Commun Biol 2023, 6:130.

7. Cao G, Xuan X, Hu J, Zhang R, Jin H, Dong H: How vascular smooth muscle cell phenotype switching contributes to vascular disease. Cell Commun Signal 2022, 20:180.

8. Miano JM, Fisher EA, Majesky MW: Fate and State of Vascular Smooth Muscle Cells in Atherosclerosis. Circulation 2021, 143:2110–2116.

9. Elmarasi M, Elmakaty I, Elsayed B, Elsayed A, Zein JA, Boudaka A, Eid AH: Phenotypic switching of vascular smooth muscle cells in atherosclerosis, hypertension, and aortic dissection. J Cell Physiol 2024, 239:e31200.

10. Shankman LS, Gomez D, Cherepanova OA, Salmon M, Alencar GF, Haskins RM, Swiatlowska P, Newman AA, Greene ES, Straub AC, et al: KLF4-dependent phenotypic modulation of smooth muscle cells has a key role in atherosclerotic plaque pathogenesis. Nat Med 2015, 21:628–637.

11. Yap C, Mieremet A, de Vries CJM, Micha D, de Waard V: Six Shades of Vascular Smooth Muscle Cells Illuminated by KLF4 (Krüppel-Like Factor 4). Arterioscler Thromb Vasc Biol 2021, 41:2693–2707.

12. Zhai X, Cao S, Wang J, Qiao B, Liu X, Hua R, Zhao M, Sun S, Han Y, Wu S, et al: Carbonylation of Runx2 at K176 by 4-Hydroxynonenal Accelerates Vascular Calcification. Circulation 2024, 149:1752–1769.

13. Yang H, Zhou T, Sorenson CM, Sheibani N, Liu B: Myeloid-Derived TSP1 (Thrombospondin-1) Contributes to Abdominal Aortic Aneurysm Through Suppressing Tissue Inhibitor of Metalloproteinases-1. Arterioscler Thromb Vasc Biol 2020, 40:e350–e366.

14. Zeng Z, Ma Y, Hu L, Tan B, Liu P, Wang Y, Xing C, Xiong Y, Du H: OmicVerse: a framework for bridging and deepening insights across bulk and single-cell sequencing. Nat Commun 2024, 15:5983.

15. Patro R, Duggal G, Love MI, Irizarry RA, Kingsford C: Salmon provides fast and bias-aware quantification of transcript expression. Nat Methods 2017, 14:417–419.

16. Browaeys R, Saelens W, Saeys Y: NicheNet: modeling intercellular communication by linking ligands to target genes. Nat Methods 2020, 17:159–162.

17. Matsumoto H, Moir LM, Oliver BG, Burgess JK, Roth M, Black JL, McParland BE: Comparison of gel contraction mediated by airway smooth muscle cells from patients with and without asthma. Thorax 2007, 62:848–854.

18. Shalhout SZ, Yang PY, Grzelak EM, Nutsch K, Shao S, Zambaldo C, Iaconelli J, Ibrahim L, Stanton C, Chadwick SR, et al: YAP-dependent proliferation by a small molecule targeting annexin A2. Nat Chem Biol 2021, 17:767–775.

19. Das A, Yang CS, Arifuzzaman S, Kim S, Kim SY, Jung KH, Lee YS, Chai YG: High-Resolution Mapping and Dynamics of the Transcriptome, Transcription Factors, and Transcription Co-Factor Networks in Classically and Alternatively Activated Macrophages. Front Immunol 2018, 9:22.

20. Deroo S, Deatrick KB, Henke PK: The vessel wall: A forgotten player in post thrombotic syndrome. Thromb Haemost 2010, 104:681–692.

21. Deatrick KB, Eliason JL, Lynch EM, Moore AJ, Dewyer NA, Varma MR, Pearce CG, Upchurch GR, Wakefield TW, Henke PK: Vein wall remodeling after deep vein thrombosis involves matrix metalloproteinases and late fibrosis in a mouse model. J Vasc Surg 2005, 42:140–148.

22. Arevalo Martinez M, Ritsvall O, Bastrup JA, Celik S, Jakobsson G, Daoud F, Winqvist C, Aspberg A, Rippe C, Maegdefessel L, et al: Vascular smooth muscle-specific YAP/TAZ deletion triggers aneurysm development in mouse aorta. JCI Insight 2023, 8.

23. Garoffolo G, Sluiter TJ, Thomas A, Piacentini L, Ruiter MS, Schiavo A, Salvi M, Saccu C, Zoli S, Chiesa M, et al: Blockade of YAP Mechanoactivation Prevents Neointima Formation and Adverse Remodeling in Arterialized Vein Grafts. J Am Heart Assoc 2025, 14:e037531.

24. Wang X, Hu G, Gao X, Wang Y, Zhang W, Harmon EY, Zhi X, Xu Z, Lennartz MR, Barroso M, et al: The induction of yes-associated protein expression after arterial injury is crucial for smooth muscle phenotypic modulation and neointima formation. Arterioscler Thromb Vasc Biol 2012, 32:2662–2669.

25. Prandoni P, Fluharty M, Schellong S, Bounameaux H, Haas S, Mantovani LG, Sholkamy S, Kondo K, Gibbs H, Jing ZC, et al: Sex differences in venous thromboembolism outcomes: findings from the GARFIELD-VTE registry. Eur J Intern Med 2025:106492.

