## Supplementary figures and images for "Inflammatory macrophages drive smooth muscle dedifferentiation via YAP signaling in murine deep vein thrombosis"

### Supplemental Material

Fig. 4D

MYH11

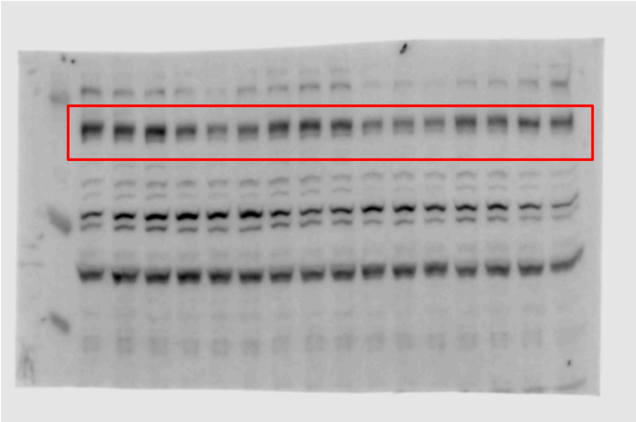

β-Actin

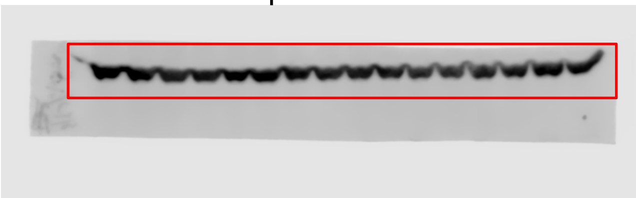

Fig. 6A

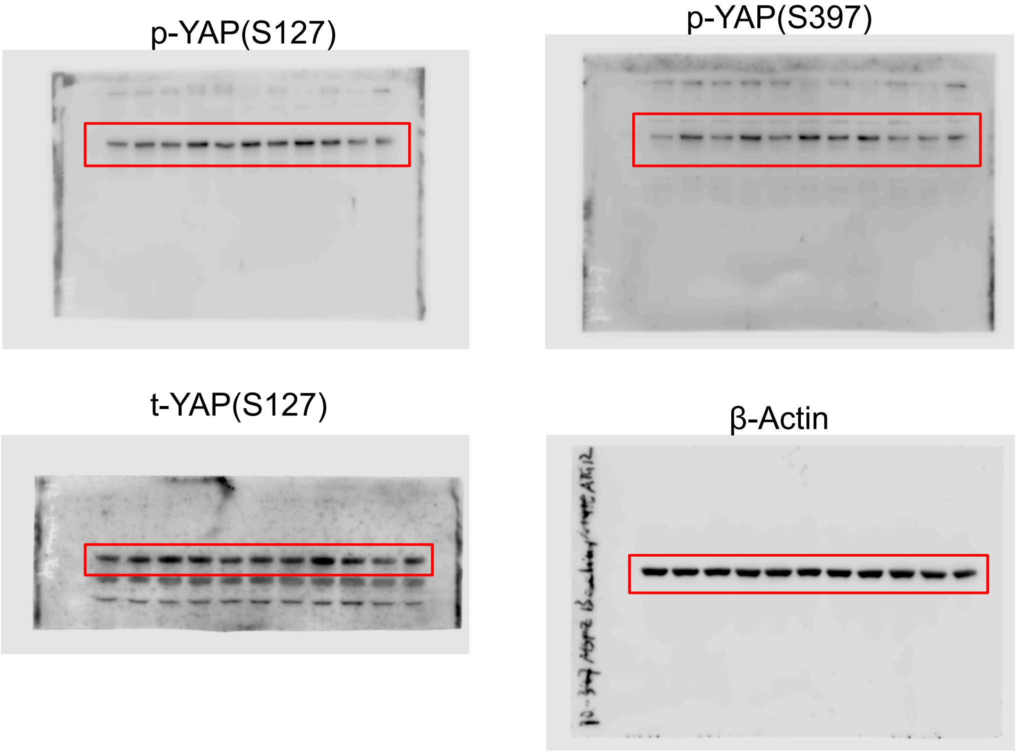


Fig. 6F

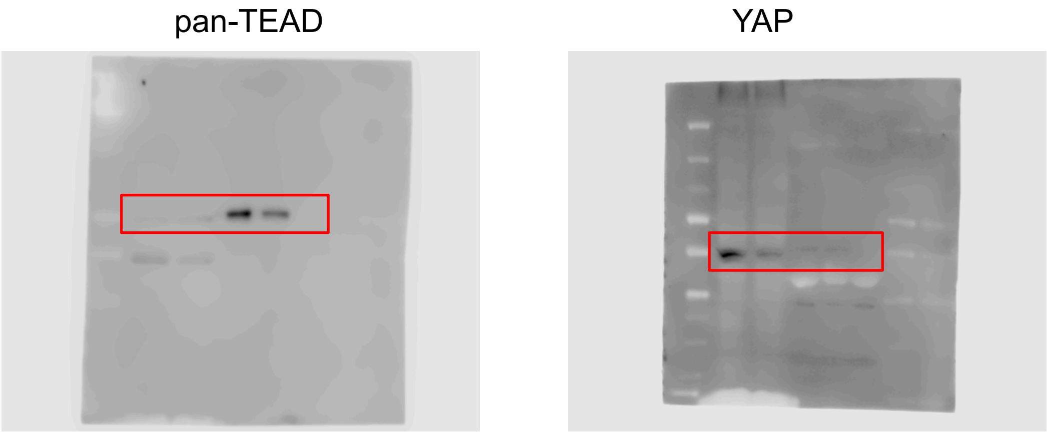

Fig. 7A

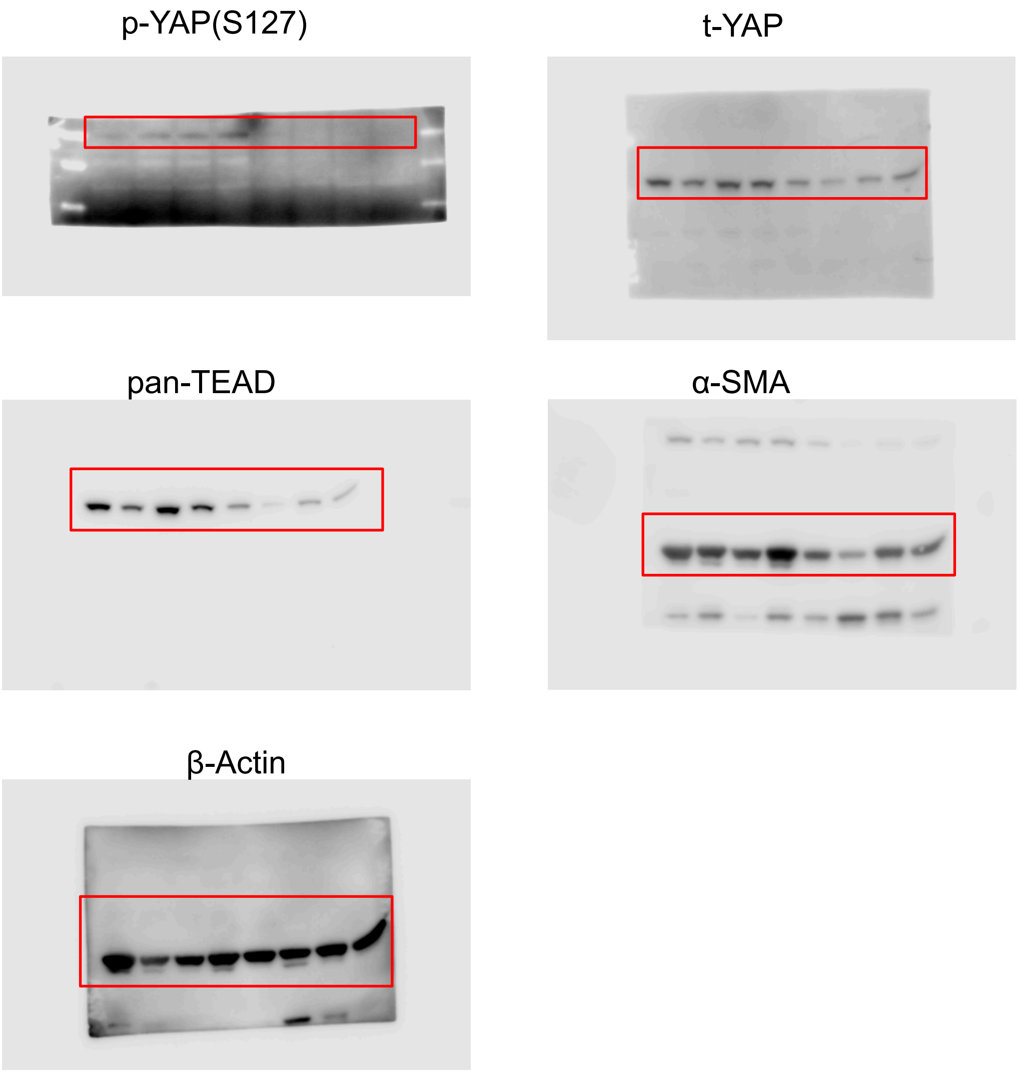

Fig. 8C

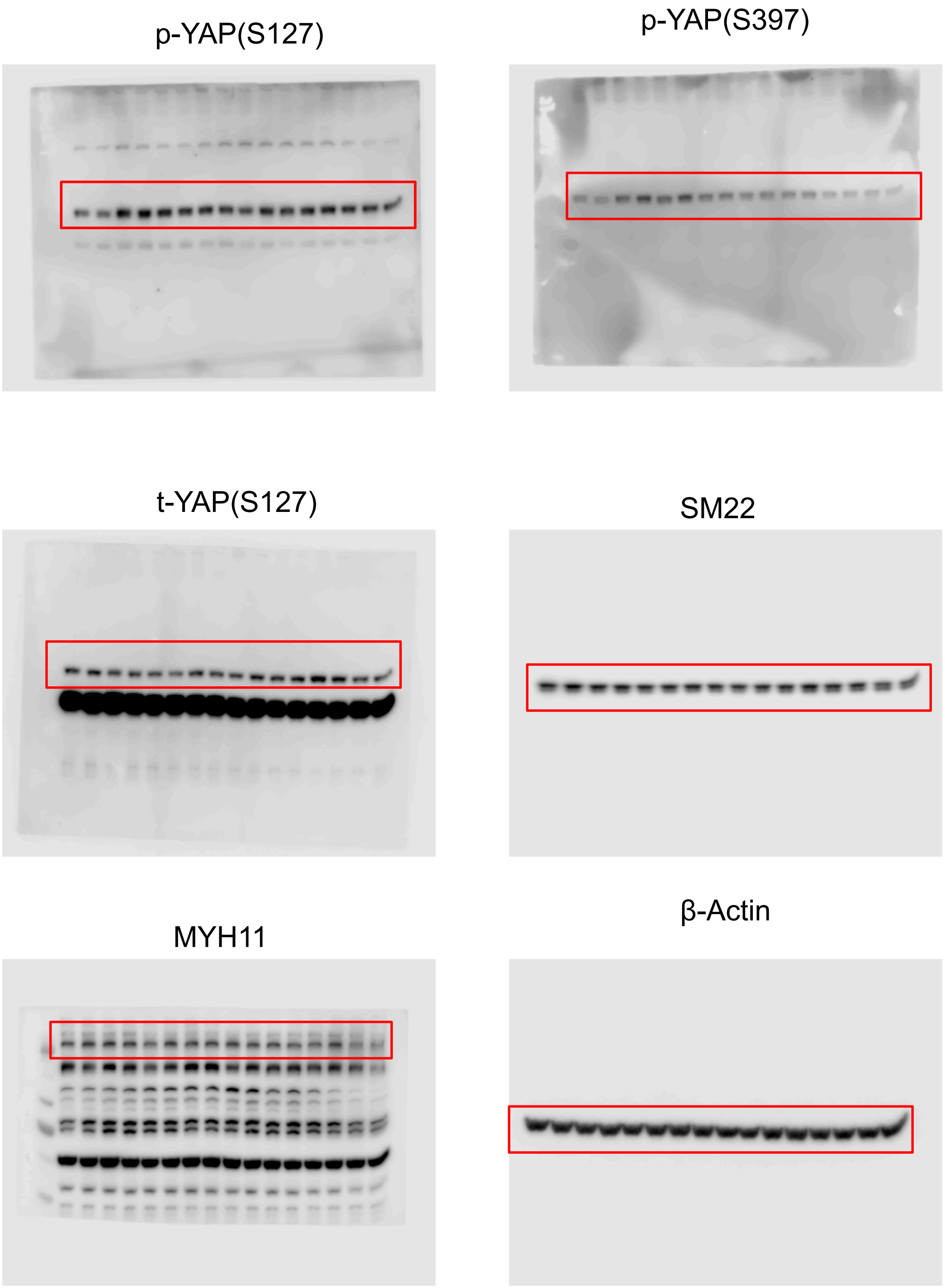
